## Supplementary Information for "The activity of soil microbial taxa in the rhizosphere predicts the success of root colonization"

Submitted to: M System

**Authors:** Jennifer E. Harris[1,2,3], Regina B. Bledsoe[1], Sohini Guha[1], Haneen Omari[3], Sharifa G. Crandall[4], Liana T. Burghardt [1,2,*], Estelle M. Couradeau[2,3,*,**]

[1] Department of Plant Science, Penn State University, State College, PA

[2] Ecology Program, Huck Institute for the Life Sciences, Penn State University, State College, PA

[3] Department of Ecosystem Science and Management, Penn State University, State College, PA

[4] Department of Plant Pathology and Environmental Microbiology, Penn State University, State College, PA

[*]equal contribution

[**] corresponding author

**Supplemental Methods:**

*LC-MS soil metabolomics:*

The natural level of methionine in soils was tested with LS-MS metabolomics. Methionine from soils was extracted by adding 5 ml of LC-MS grade water to 1g of soil in 3 replicates. Samples were incubated for 1 hour at 4C and vortexed for 5s every 15 minutes. Samples were centrifuged at 3220xg for 15 minutes at 4C. We filtered the supernatant with a 0.45 µm filter and freeze-dried with Freezone 12L -50 Lyophilizer (Labcono) for 72 hours. Samples (5 uL) were separated by reverse phase HPLC using a Prominence 20 UFLCXR system (Shimadzu), with a Waters BEH C18 column (100mm × 2.1mm 1.7 um particle size) maintained at 55°C and a 20-minute aqueous acetonitrile gradient, at a flow rate of 250 uL/min. Solvent A was HPLC-grade water with 0.1% formic acid, and Solvent B was HPLC-grade acetonitrile with 0.1% formic acid. The initial conditions were 97% A and 3% B, increasing to 45% B at 10 min and 75% B at 12 min, where it was held at 75% B until 17.5 min before returning to the initial conditions. The eluate was delivered into a 5600 (QTOF) TripleTOF using a Duospray™ ion source (all Sciex). The capillary voltage was set at 5.5 kV in positive ion mode with a declustering potential of 80V. The mass spectrometer was operated with a 100 ms TOF scan from 50 to 500 m/z and 16 MS/MS product ion scans (100 ms) per duty cycle using a collision energy of 50V with a 20V spread. The Penn State Huck Metabolomics core quantified methionine peaks in Sciex Analyst software.

*Probing Microbial Activity with BONCAT:*

We used BONCAT to label metabolically active cells in the bulk soil, rhizosphere, root endosphere, and nodules. After incubation with HPG and cell extraction, a copper-catalyzed click reaction was performed to attach FAM picolyl azide dye (Ex/Em 490/510 nm) to the newly-made proteins during the greenhouse incubation. We followed the protocol described in Couradeau et al. 2019 (full detailed protocol in supplemental methods. ( Briefly, 700 µl of each cell extract was captured on a 0.22 µm Millipore isopore filter, and the liquid was removed by vacuum filtration. Then, the sample-containing filter was transferred to a glass microscope slide, where we added 80 µl of the reaction mix. The reaction mix consisted of 5 µl copper sulfate ($CuSO4$ 100 µM final concentration), of 10 µl tris-hydroxypropyltriazolylmethylamine (THPTA, 500 µM final concentration), and of

3.3 μl (FAM picolyl azide dye, 5 μM final concentration), buffered in of 50 μl of 5 mM sodium ascorbate freshly prepared in 1M PBS and 50 μl of 5 mM aminoguanidine HCl freshly prepared in 1M PBS and 880 μl of 1M PBS. All reagents were purchased from Click Chemistry Tools. The sample-containing filter was sealed with a cover slip and incubated in a dark place at room temperature for 30 minutes. After the incubation, the excess dye was washed off by soaking the sample PBS for five minutes three times. Cells were removed from the filter by vortexing in PBS-tween for 5 min. Finally, we added SYTO59 DNA counterstain (Invitrogen, Ex/Em 622/645 nm) at a final concentration of 0.5 μM.

*Flow cytometry gating and sorting:*

The evaluation of BONCAT fluorescence was first performed on a BD LSRFortessa Cell Analyzer (Becton, Dickinson, and Company). The threshold size was set to above 0.2 μm determined by Flow Cytometry Sub-Micron Particle Size Reference Kit Beads (Invitrogen). BD LSRFortessa Cell Analyzer was set up to capture the FAM picolyl azide dye (Ex/Em 490/510 nm) in the green channel of a 488 nm blue laser and the SYTO59 DNA counterstain (Ex/Em 622/645 nm) in the red channel of a 630 nm red laser. Viable cells were identified by drawing a SYTO59-positive gate against an unstained control. Active cells were gated from the SYTO59-positive events using a nested gating strategy. Active cells were determined by BONCAT FAM picolyl azide-positive events against a water-incubated clicked control to account for the background fluorescence from the click reaction. The false positivity recovery rate was capped at 0.2% in the gating process. We used the same gating strategy for Fluorescence-activated cell sorting (FACS) on a MoFlo Astrios Cell Sorter (Beckman Cloutier) to sort cells from plant compartments (rhizosphere, root, nodule). We could not recover enough active cells from the bulk soil samples due to their low cell event numbers and low activity to include them in the downstream process. We sorted from each sample (mean= 82,000) for the active cell fraction, and only samples with at least 50k Active cells were included in our analysis (see Supplementary Table 2 for cells sorted counts from each sample). For the viable cell fraction, we sorted 100k Viable cells from each sample, using unclicked cell extract and directly labeled with a final concentration of 0.5 μM of SYBR Green (Invitrogen, Ex/Em 498/522 nm) as a general DNA probe.

*DNA extractions and 16S rRNA amplicon sequencing:*

We sequenced five plants' active and viable fractions from three compartments (rhizosphere, root endosphere, and nodule). We also sequenced Total DNA from Bulk Soil and rhizosphere (four replicates). For active and viable fractions, our library prep protocol was adapted from Reichart et al. 2020 [32]. Cells were centrifuged at 14000g for 10 min at 4°C. The supernatant was pipetted out, and 9.2 µl of nuclease-free water was added. Cells were then lysed with prepGEM® Bacteria Kit cell lysing kit following manufacturer instructions (New England Biolabs). We extracted total DNA from all rhizosphere and bulk soil samples with the DNeasy PowerSoil Pro Kit (Qiagen). We used 515f-Y (GTGYCAGCMGCCGCGGTAA, Parada 2016) and 806R-B GGACTACNVGGGTWTCTAAT, April 2015) primers to amplify the 16S rRNA gene from lysed cells pellets and total DNA. Our final PCR reaction mixture included a final concentration of 1x Invitrogen Platinum Super Master Mix and .2 µM of each primer. The thermocycler program included initial denaturation of 98C for 2 minutes, 30X cycles of 98°C for 10 seconds, 56.2°C for 20 seconds, 72°C for 15 seconds, and a final elongation of 72°C for 5 minutes. The Huck Life Science Genomics Core carried out library preparation and indexing. Of the 30 samples (2 fractions X 3 compartments X 5 reps) we sorted cells from, 27 samples had successful library preparation, resulting in at least four complete reps for each treatment. The successful libraries were sequenced on an Illumina MiSeq platform using 250 bp x 250 bp paired-end sequencing and aiming at 100,000 reads per sample. The sequencing run produced 7,187,623 reads, averaging 107,255 paired reads per sample.

*Microscopy:*

We used confocal microscopy to visualize BONCAT-labeled active microbial cells in the nodule. Pre-inoculated *Trifolium incarnatum* was grown in potting soil in the greenhouse. After eight weeks, three plants were dosed with HPG dissolved in water at a final concentration of 0.038 µmol HPG per g soil, and one plant with HPG-free water as a negative control. After 24 hours, nodules were picked, washed, and stored in PIPES buffer for ~16 hours. Next, nodules were embedded nodules in Epredia Cryomatrix resin and frozen. 50 µm thick sections were cut from distal to proximal end onto poly-lysine-coated slides with a Leica CM1950 Cryostat. After sectioning, the cryomatrix was dissolved by soaking sections in DI water for 5 minutes. The BONCAT reaction mix was the same as our flow cytometry samples. However, we substituted Picolyl-Azide-5/6-FAM for Cy3 Picolyl Azide (Click Chemistry Tools, Ex/Em 553/568) because plant cell walls showed the least autofluorescence in this Ex/Em range. 100 µl of BONCAT reaction mix was added to each section and incubated for 1 hour at room temperature in the dark. After incubation, excess dye was removed by washing with PBS three times. BONCAT labeled sections were imaged on the Leica SP8 DIVE multiphoton microscope. BONCAT Cy3 dye was excited on a diode 553-laser line and captured fluorescence with sensitive hybrid detectors (HyDs). For visualizing the general nodule structure, sections were incubated for 5 minutes with .1% toluidine blue. Toluidine blue images were captured on a Zeiss Axio Observer fluorescence microscope with an Axio 208 color camera. Images were processed using ImageJ (Fiji) [31]. Using BONCAT and toluidine blue, we could identify zones of nodules with metabolically active microbial cells.

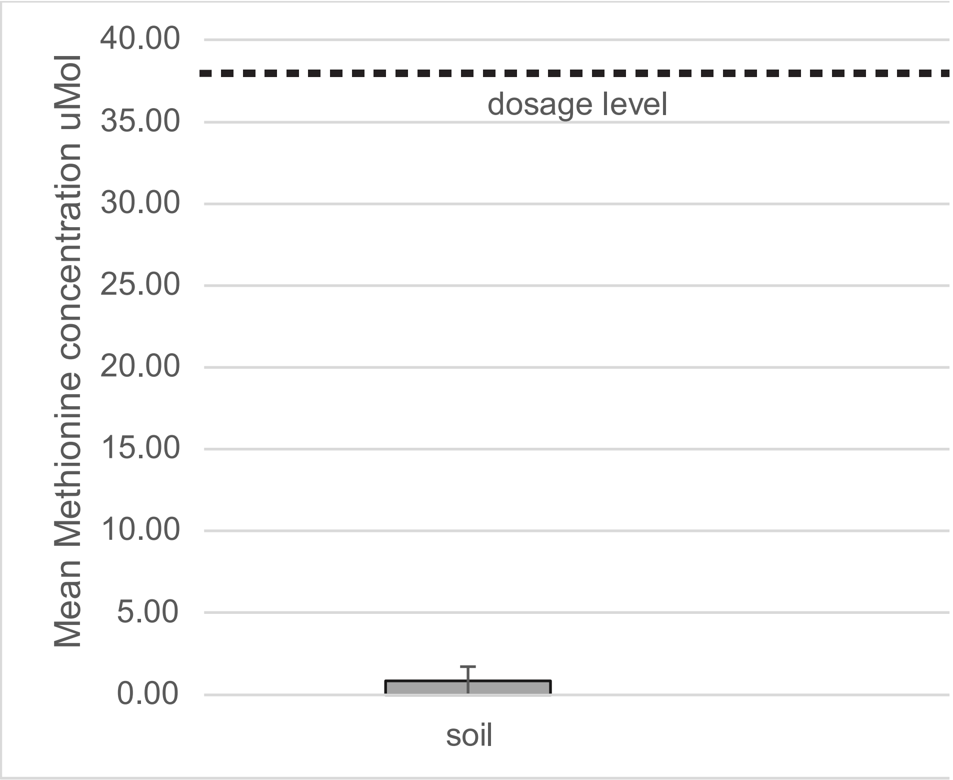


**Supplementary Figure 1:** LC-MS analysis of soil extract shows that natural methionine in our soils was lower than dosage levels. Data are mean methionine concentration in uM (n= 2, one sample was below the detection limit). Concetrations are calculated from a standard curve of know concentrations of Methionine. Methionine Peak area adjusted by the area of an internal standard (Chlorpropamide). Error bars are standard deviation.

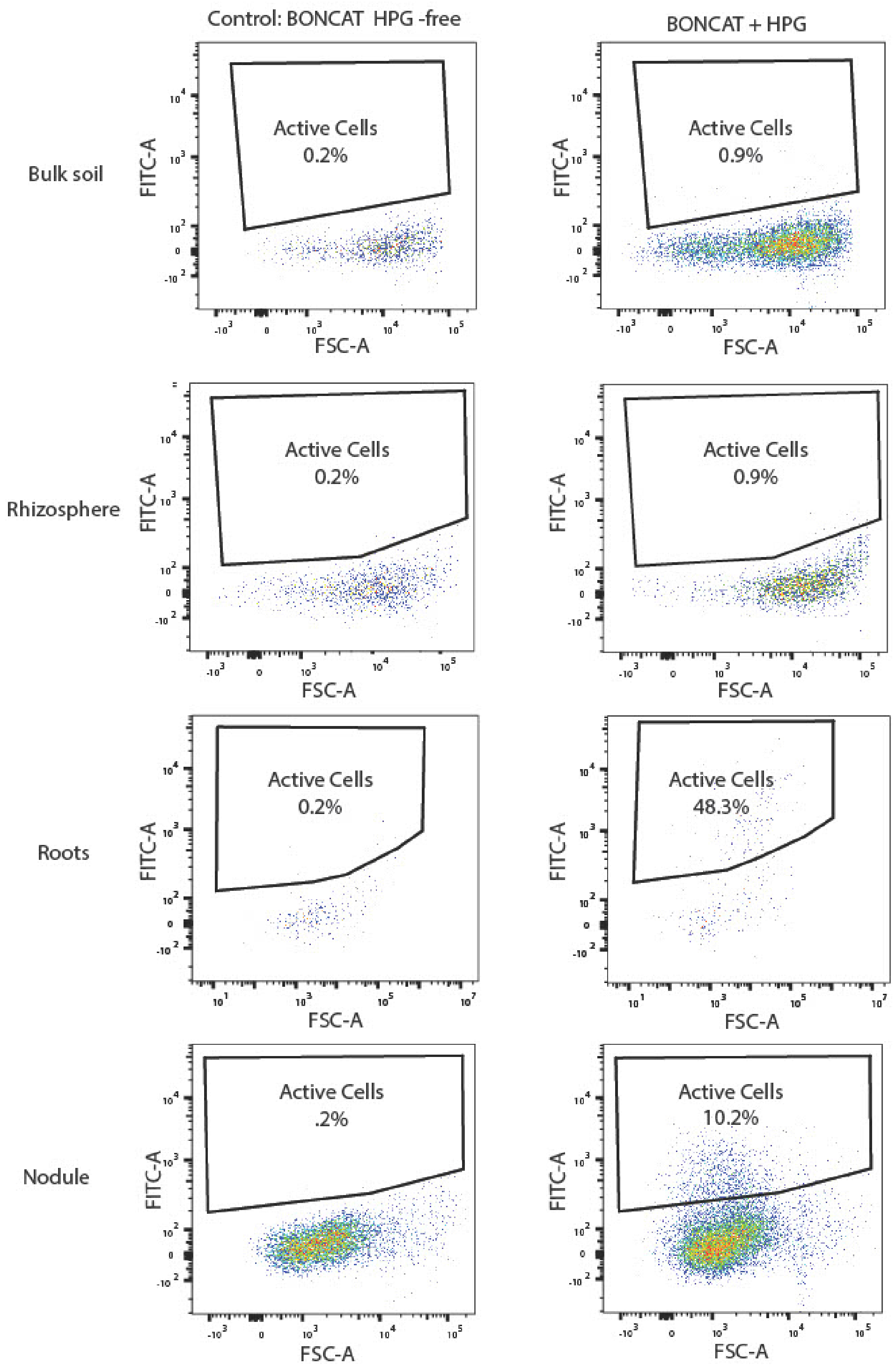


**Supplementary Figure 2:** Flow Cytometry of BONCAT labeled cells in bulk soil, rhizosphere, roots, and nodules. All events shown were SYTO positive. Flow cytometer events are plotted according to their forward scatter signal (FSC, x-axis) and BONCAT fluorescence (FITC, y-axis) in a log-log scale. Increased cell event density is indicated by color. The BONCAT+ active cells gate is displayed as a box in each plot, and the percent cells in the BONCAT+ gate are indicated in the box. The left column shows water-incubated controls given the BONCAT reaction mix and clicked. The right column shows samples incubated with HPG, given with BONCAT reaction, and clicked.

**Supplementary Table 1:** Likelihood ratio test results of the proportion of active cells in a quasibinomial model(proportion active ~ fraction + rep). Data was collected on the BD Fortessa flow cytometer.

| *Comparison* | *df* | *Deviance* | *Resid.df* | *Resid. Deviance* | *P value* | |
|---|---|---|---|---|---|---|
| Bulk soil vs Rhizosphere | 1 | 16.3 | 12 | 95.7 | 0.15 | |
| Bulk soil vs Roots | 1 | 3635 | 12 | 2016 | 3.30E-05 | * |
| Bulk soil vs Nodule | 1 | 3953 | 12 | 1895 | 1.67E-05 | * |
| Rhizosphere vs Roots | 1 | 2997 | 12 | 1986 | 1.57E-04 | * |
| Rhizosphere vs Nodule | 1 | 3125 | 12 | 1865 | 1.24E-04 | * |
| Roots vs Nodule | 1 | 12.2 | 12 | 3785 | 0.86 | |

*Significant in the model (p<.05)

**Supplementary Table 2:** 16S RNA samples description, numbers of cells sorted, and reads passing quality control, number of unrarefied ASVs, and number of ASVs when rarefied to 40592 reads.

| *Sample ID* | *Compartment* | *Fraction* | *Replicate Pot* | *No. Cells sorted* | *Raw Reads* | *Reads passing QC* | *No. ASVS* | *No. ASVS rarefied* |
|---|---|---|---|---|---|---|---|---|
| *C10N-POS_S65* | Nodule | Active | 1 | 60732 | 159315 | 138515 | 24 | 23 |
| *C1N-POS_S61* | Nodule | Active | 2 | 70966 | 153712 | 92476 | 25 | 24 |
| *C2N-POS_S62* | Nodule | Active | 3 | 66599 | 173557 | 135988 | 26 | 23 |
| *C5N-POS_S63* | Nodule | Active | 4 | 100000 | 188756 | 130178 | 22 | 22 |
| *C7N-POS_S64* | Nodule | Active | 5 | 100000 | 138863 | 100742 | 30 | 29 |
| *C10R-POS_S30* | Rhizosphere | Active | 1 | 97,888 | 101997 | 83847 | 864 | 855 |
| *C1R-POS_S27* | Rhizosphere | Active | 2 | 100000 | 105336 | 80724 | 1091 | 1086 |
| *C2R-POS_S28* | Rhizosphere | Active | 3 | 100000 | 91265 | 67920 | 1017 | 1014 |
| *C5R-POS_S29* | Rhizosphere | Active | 4 | 100000 | 88102 | 61804 | 1007 | 1005 |
| *C10E-POS_S60* | Root | Active | 1 | 50506 | 147464 | 85169 | 116 | 114 |
| *C1E-POS_S31* | Root | Active | 2 | 58919 | 145971 | 82832 | 152 | 147 |
| *C2E-POS_S32* | Root | Active | 3 | 73549 | 154251 | 76644 | 112 | 109 |
| *C5E-POS_S33* | Root | Active | 4 | 100000 | 175034 | 100352 | 120 | 114 |
| *C7E-POS_S34* | Root | Active | 5 | 21364* | 227295 | 142404 | 150 | 141 |
| *C10N-SYBR_S26* | Nodule | Viable Cell | 1 | 100000 | 188796 | 170390 | 49 | 47 |
| *C1N-SYBR_S13* | Nodule | Viable Cell | 2 | 100000 | 234920 | 180861 | 42 | 37 |
| *C2N-SYBR_S15* | Nodule | Viable Cell | 3 | 100000 | 125690 | 114107 | 26 | 24 |
| *C7N-SYBR_S25* | Nodule | Viable Cell | 5 | 100000 | 153058 | 130912 | 22 | 21 |
| *C10R-SYBR_S20* | Rhizosphere | Viable Cell | 1 | 100000 | 78509 | 48992 | 1137 | 1137 |
| *C1R-SYBR_S16* | Rhizosphere | Viable Cell | 2 | 100000 | 121003 | 85773 | 1329 | 1320 |
| *C2R-SYBR_S17* | Rhizosphere | Viable Cell | 3 | 100000 | 126308 | 87086 | 1317 | 1309 |
| *C5R-SYBR_S18* | Rhizosphere | Viable Cell | 4 | 100000 | 103848 | 68636 | 1417 | 1410 |
| *C7R-SYBR_S19* | Rhizosphere | Viable Cell | 5 | 100000 | 95994 | 62332 | 1196 | 1194 |
| *C1E-SYBR_S21* | Root | Viable Cell | 2 | 100000 | 157198 | 105567 | 100 | 97 |
| *C2E-SYBR_S22* | Root | Viable Cell | 3 | 100000 | 154515 | 103885 | 76 | 75 |
| *C5E-SYBR_S23* | Root | Viable Cell | 4 | 100000 | 161291 | 117193 | 58 | 54 |
| *C7E-SYBR_S24* | Root | Viable Cell | 5 | 100000 | 217885 | 193245 | 54 | 46 |
| *C10B-DNA_S4* | Bulk Soil | Total DNA | 1 | NA | 82320 | 40592 | 1342 | 1342 |
| *C2B-DNA_S1* | Bulk Soil | Total DNA | 3 | NA | 103445 | 46386 | 1454 | 1452 |
| *C5B-DNA_S2* | Bulk Soil | Total DNA | 4 | NA | 109593 | 59049 | 1865 | 1859 |
| *C7B-DNA_S3* | Bulk Soil | Total DNA | 5 | NA | 90895 | 49548 | 1474 | 1472 |
| *S10-DNA_S9* | Bulk Soil | Total DNA | 6 | NA | 107278 | 59880 | 1955 | 1945 |
| *S2-DNA_S5* | Bulk Soil | Total DNA | 7 | NA | 80802 | 48575 | 1551 | 1549 |
| *S3-DNA_S6* | Bulk Soil | Total DNA | 8 | NA | 92770 | 49128 | 1645 | 1640 |
| *S8-DNA_S7* | Bulk Soil | Total DNA | 9 | NA | 83071 | 50225 | 1539 | 1537 |
| *C10R-DNA_S12* | Rhizosphere | Total DNA | 1 | NA | 112015 | 58573 | 1430 | 1413 |
| *C1R-DNA_S8* | Rhizosphere | Total DNA | 2 | NA | 101059 | 50926 | 1294 | 1292 |
| *C2R-DNA_S10* | Rhizosphere | Total DNA | 3 | NA | 114438 | 62767 | 1533 | 1514 |
| *C5R-DNA_S11* | Rhizosphere | Total DNA | 4 | NA | 103825 | 67223 | 1534 | 1521 |
| *C7R-DNA_S14* | Rhizosphere | Total DNA | 5 | NA | 123393 | 67185 | 1507 | 1484 |
| *CTL_S66* | Control | Flow Cyto | NA | NA | 104767 | 63774 | 1642 | 1640 |
| *BEADS* | Control | PCR | NA | NA | 92886 | 72911 | 815 | 812 |

*Data point omitted from analysis because of low number of cells

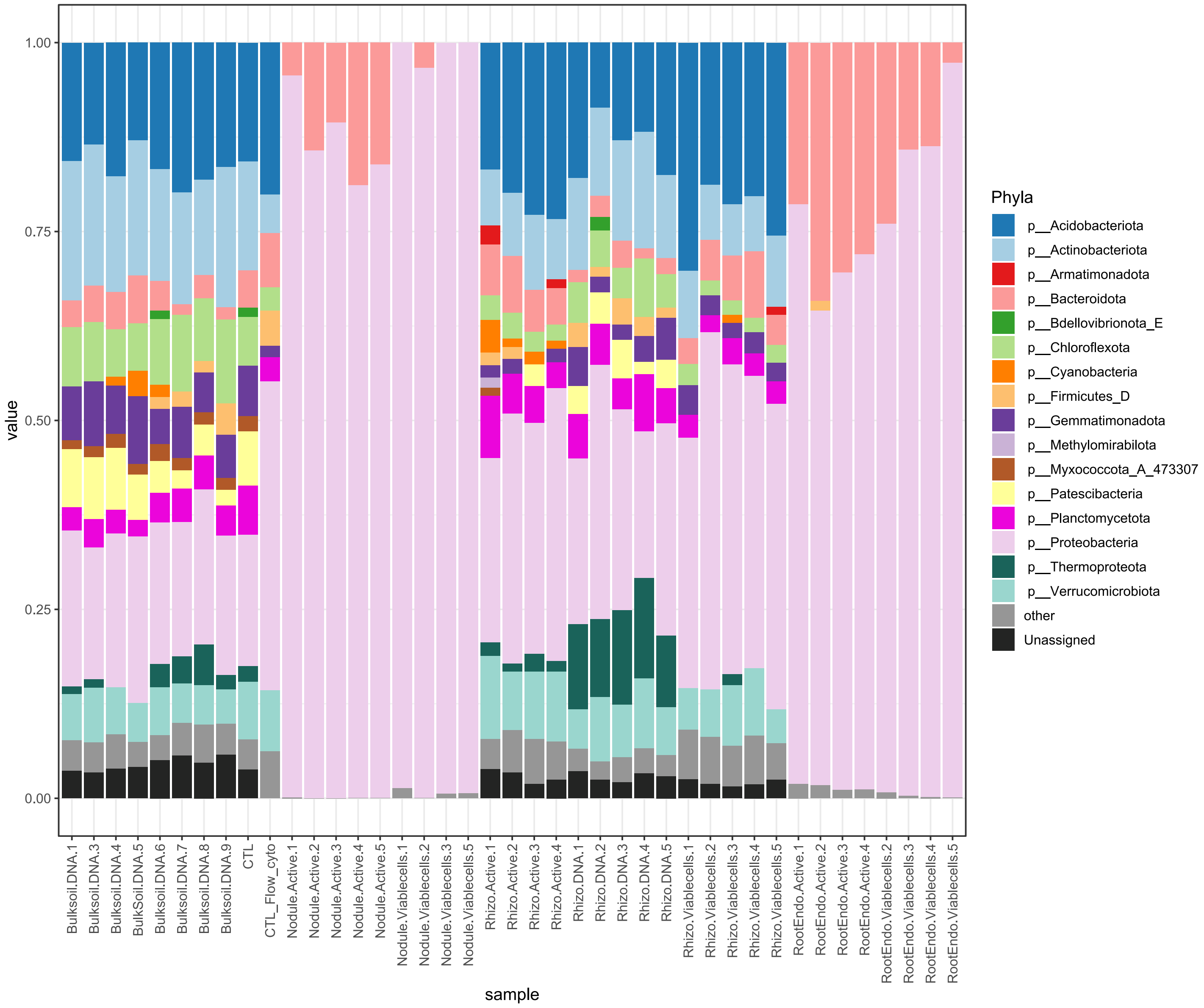


**Supplementary Figure 3:** Phyla level microbial community composition across all samples. Each stacked bar is a sample. Sample names are on the X-axis. Taxa with less than 1% abundance were placed in "other." ASVs are unrarefied.

**Supplementary Table 3:** ANOVA comparisons for Shannon diversity of microbial communities. We subset data into fractions to compare Shannon diversity across compartments (Shannon diversity ~ compartment). We subset data into each compartment to compare the diversity between active and viable fractions. Shannon diversity is calculated on rarefied reads in R package phyloseq.

| *Comparison*[1] | *df* | *Effect Size* | *Standard Error* | *t value* | *p-value* | |
|---|---|---|---|---|---|---|
| Total DNA: Bulk Soil vs. Rhizosphere | 7 | -0.30 | 0.10 | -3.1 | 0.018 | * |
| Viable Cells: Rhizosphere vs. Roots | 7 | -5.06 | 0.16 | -31.1 | 9.1E-09 | * |
| Viable Cells: Roots vs. Nodule | 6 | -0.26 | 0.19 | -1.4 | 0.212 | |
| Active Cells: Rhizosphere vs. Roots | 6 | -4.44 | 0.11 | -40.2 | 1.6E-8 | * |
| Active Cells: Roots vs. Nodule | 7 | -0.60 | 0.12 | -5.2 | 1.3E-03 | * |
| Rhizosphere: Total DNA vs. Viable Cells | 8 | -0.20 | 0.10 | -1.9 | 0.082 | |
| Rhizosphere: Total DNA vs. Active Cells | 7 | -0.33 | 0.10 | -3.3 | 0.012 | * |
| Rhizosphere: Active vs. Viable cells | 7 | 0.13 | 0.10 | 1.3 | 0.248 | |
| Roots: Active vs. Viable Cells | 6 | -0.49 | 0.18 | -2.7 | 0.037 | * |
| Nodule: Active vs. Viable Cells | 7 | -0.15 | 0.12 | -1.2 | 0.263 | |

*Significant in the model (p<.05)
[1]Reference level/group listed first

**Supplementary Table 4:** ANOVA comparisons for the number of ASVs microbial communities. We subset into fractions to compare No. Of ASVs detected across compartments (No. ASVs ~ compartment). To compare the diversity between active and viable fractions, we subset data into each compartment. No. of ASVs from rarefied reads.

| *Comparison*[1] | *df* | *Effect Size* | *Standard Error* | *t value* | *p-value* | |
|---|---|---|---|---|---|---|
| Total DNA: Bulk Soil vs. Rhizosphere | 7 | -86 | 110 | -0.8 | 0.457 | |
| Viable Cells: Rhizosphere vs. Roots | 7 | -1206 | 55 | -21.6 | 1.2E-07 | * |
| Viable Cells: Roots vs. Nodule | 6 | -36 | 13 | -2.8 | 0.033 | * |
| Active Cells: Rhizosphere vs. Roots | 7 | -965 | 43 | -22.7 | 8.3E-08 | * |
| Active Cells: Roots vs. Nodule | 7 | -97 | 8 | -12.4 | 5.1E-06 | * |
| Rhizosphere: Total DNA vs. Viable Cells | 8 | -170.8 | 65 | -2.6 | 0.030 | * |
| Rhizosphere: Total DNA vs. Active Cells | 7 | -455 | 65 | -7.0 | 2.0E-04 | * |
| Rhizosphere: Active vs. Viable cells | 7 | 284 | 70 | 4.1 | 4.7E-03 | * |
| Roots: Active vs. Viable Cells | 6 | -53 | 14 | -3.7 | 0.010 | * |
| Nodule: Active vs. Viable Cells | 7 | 8 | 6 | 1.5 | 0.185 | |

*Significant in the model (p<.05)
[1]Reference level/group listed first

**Supplementary Table 5:** ANOVA comparisons for evenness microbial communities. We subset into fractions to compare Evenness across compartments. To compare the diversity between active and viable fractions, we subset data into each compartment. Evenness is calculated using rarefied reads. Evenness calculated as Evenness = Shannon diversity/ (Natural log(Observed ASVs))

| *Comparison*[1] | *df* | *Effect Size* | *Standard Error* | *t value* | *p-value* | |
|---|---|---|---|---|---|---|
| Total DNA: Bulk Soil vs. Rhizosphere | 7 | -0.01 | 0.01 | -4.5 | 2.7E-03 | * |
| Viable Cells: Rhizosphere vs. Roots | 7 | -0.60 | 0.03 | -21.9 | 1.1E-07 | * |
| Viable Cells: Roots vs. Nodule | 6 | -0.01 | 0.04 | -0.3 | 0.788 | |
| Active Cells: Rhizosphere vs. Roots | 7 | -0.54 | 0.02 | -30.8 | 7.8E-08 | * |
| Active Cells: Roots vs. Nodule | 7 | -0.02 | 0.03 | -0.6 | 0.569 | |
| Rhizosphere: Total DNA vs. Viable Cells | 8 | -0.01 | 0.01 | -1.0 | 0.353 | |
| Rhizosphere: Total DNA vs. Active Cells | 7 | 7E-04 | 0.01 | 0.1 | 0.939 | |
| Rhizosphere: Active vs. Viable cells | 7 | -0.01 | 0.01 | -1.0 | 0.348 | |
| Roots: Active vs. Viable Cells | 6 | -0.07 | 0.03 | -2.2 | 0.073 | |
| Nodule: Active vs. Viable Cells | 7 | -0.06 | 0.04 | -1.7 | .131 | |

*Significant in the model (p<.05)
[1]Reference level/group listed first

**Supplementary Table 6:** Permutation test results of groups beta dispersion. Beta dispersion was calculated on groups' Bray Curtis dissimilarity matrixes. We did a Permutation Test of the groups' beta dispersion to determine if there was nonhomogeneity of variance. Homogeneity of variance between groups is required for a PERMANOVA.

| *Comparison* | *Df total* | *Sum sq total* | *Mean sq total* | *F value* | *No. of Permutations* | *p Value* | |
|---|---|---|---|---|---|---|---|
| Full community Between Compartments | 38 | 0.737 | 0.214 | 71.02 | 999 | 0.001 | * |
| Bulk soil + Rhizosphere: Between Compartments | 21 | 0.063 | 0.026 | 12.55 | 999 | 0.001 | * |
| Rhizosphere: All fractions | 13 | 0.040 | 0.017 | 29.28 | 999 | 0.001 | * |
| Rhizosphere: Active vs Viable Cells | 8 | 0.018 | 0.014 | 19.80 | 999 | 0.001 | * |
| Total DNA: Bulk Soil vs Rhizosphere | 8 | 0.004 | 0.001 | 2.30 | 999 | 0.166 | |
| Roots + Nodule: Between Compartments | 16 | 0.075 | 0.014 | 2.27 | 999 | 0.131 | |
| Roots + Nodule: Active vs. Viable Cells | 16 | 0.108 | 0.010 | 0.43 | 999 | 0.511 | |
| Roots: Active vs. Viable Cells | 7 | 0.031 | 0.005 | 0.05 | 999 | 0.884 | |
| Nodule: Active vs. Viable Cells | 8 | 0.023 | 0.004 | 0.30 | 999 | 0.592 | |

*Significant value (p<.05) indicates nonhomogeneous variance between groups.

**Supplementary Table 7:** PERMANOVA test results calculated with Bray Curtis distances. Full data set was subset to test for differences between comparisons.

| *Comparison* | *Df total* | *Sum Sq total* | *$R^2$* | *F value* | *p-value* | |
|---|---|---|---|---|---|---|
| Bulk soil + Rhizosphere: Comparing Total DNA fractions | 21 | 1.20 | 0.26 | 7.14 | 0.001 | * |
| Roots + Nodule: Between Compartments | 16 | 0.60 | 0.36 | 8.26 | 0.008 | * |
| Roots + Nodule: Between Fractions | 16 | 0.60 | 0.29 | 6.16 | 0.028 | * |
| Roots: Between Fractions | 7 | 0.26 | 0.60 | 8.92 | 0.032 | * |
| Nodule: Between Fractions | 8 | 0.13 | 0.51 | 7.30 | 0.028 | * |

*Significant differences between groups (p<.05)

**Supplementary Table 8:** Differentially abundant taxa in the rhizosphere. Taxa shown met the criteria for differentially abundant in ANCOM. Log fold change reference is the Viable community. A positive Log fold change indicated ASVs enriched in the Active Community. Column name abbreviations are as follows: Rhizo total – Mean Number of reads in Rhizosphere total cell community; Rhizo active abund. – Mean Abundance in Rhizosphere BONCAT active community mean; LFC log fold change total to active community; Std. error- Standard error of the log fold change. We reported the lowest taxonomic classification identified with the greengenes2 database. Differentially abundant taxa needed to be present in at least 3 samples of in the active or the viable fraction with at least 50 reads in the rhizosphere.

| *ASV* | *Phyla* | *Lowest taxonomic classification* | *LFC* | *Std Error* | *Wilcoxon test statistic* | *Viable* | *Active* | *Adj P value* |
|---|---|---|---|---|---|---|---|---|
| *f18c54e493d5d5ad534b9c3100216416* | Proteobacteria | *Rhizobium* | 1.7 | 0.3 | -5.9 | 2600 | 2488 | 1.7E-06 |
| *2acee427328d2e8641b8738258b15927* | Proteobacteria | *Pseudomonas* | 2.3 | 0.3 | -6.7 | 758 | 1240 | 1.0E-08 |
| *509fca5a90d0b1dbec313e11c762a7e1* | Proteobacteria | *Rhizobium* | 1.7 | 0.3 | -5.7 | 384 | 362 | 8.7E-06 |
| *71582281edf705eb59a48c1a50e59a86* | Proteobacteria | *Devosia* | 1.6 | 0.3 | -5.5 | 322 | 339 | 1.8E-05 |
| *9ca5c6b3923a29ddca908fc1a47ffca1* | Proteobacteria | *Arenimonas oryziterrae* | 1.4 | 0.2 | -5.9 | 229 | 175 | 2.2E-06 |
| *ff8123ece2a6aff4b7f2750a928e83ec* | Actinobacteriota | *Acidimicrobiales* | 2.3 | 0.3 | -7.3 | 210 | 421 | 1.6E-10 |
| *757d2e39206866545010c4afcc44f748* | Actinobacteriota | *Micrococcaceae* | 2.3 | 0.3 | -7.7 | 198 | 362 | 8.4E-12 |
| *7d2d1b7d4e351f67f982eff3c2fe17cd* | Verrucomicrobiota | *Chthoniobacteraceae* | 2.6 | 0.4 | -6.6 | 188 | 533 | 1.8E-08 |
| *1b86f9be5bce7a20bcf239e2b9d628b3* | Proteobacteria | *Caulobacter* | 1.6 | 0.3 | -6.1 | 156 | 146 | 7.2E-07 |
| *3585021835f92062e3905fda1086a40f* | Proteobacteria | *Caulobacter* | 1.7 | 0.2 | -7.1 | 155 | 179 | 6.2E-10 |
| *7774b77bbabdd9fdb09f14add8d48a0f* | Acidobacteriota | *Vicinamibacterales* | 1.4 | 0.2 | -7.2 | 154 | 121 | 3.0E-10 |
| *8ad25d9768193b89b090ec9114491d2b* | Acidobacteriota | *Pyrinomonadaceae* | 1.3 | 0.3 | -4.8 | 146 | 106 | 7.2E-04 |
| *9f9aec9e6ea1a095f866295f88499642* | Acidobacteriota | *Vicinamibacterales* | 1.2 | 0.2 | -5.2 | 137 | 81 | 1.0E-04 |
| *56acd54962aaed6383d2cc96c94dce56* | Proteobacteria | *Rhizobium* | 2.5 | 0.3 | -8.8 | 135 | 304 | 7.3E-16 |
| *8b10efb60e9ddd2cb7fcd09c8fa35166* | Actinobacteriota | *Solirubrobacteraceae* | 1.8 | 0.2 | -10.2 | 122 | 145 | 1.2E-21 |
| *f6a7e105657a2c8e57f686ad1f5366f4* | Acidobacteriota | *Vicinamibacterales* | 1.7 | 0.3 | -5.1 | 122 | 140 | 1.7E-04 |
| *6c178cce4595cdbd601c6a678885170e* | Proteobacteria | *Phenylobacterium kunshanense* | 1.6 | 0.2 | -7.4 | 121 | 120 | 7.9E-11 |
| *c09563e31635d061641ff75dc777f7a9* | Proteobacteria | *Bradyrhizobium* | 1.7 | 0.3 | -5.4 | 99 | 114 | 3.1E-05 |
| *05e8b422087270ed4fcfaf95cd8b94a9* | Acidobacteriota | *Vicinamibacterales* | 1.8 | 0.3 | -6.8 | 95 | 106 | 5.2E-09 |
| *d00bdb7a37402d864b086dd2a0e7d12d* | Proteobacteria | *Ramlibacter* | 1.5 | 0.3 | -5.9 | 69 | 63 | 2.1E-06 |
| *cf2a625547909cab32e5e208dd29cdad* | Proteobacteria | *Nordella* | 1.6 | 0.3 | -5.1 | 68 | 82 | 1.5E-04 |
| *be42831588caa14346a2902a0790f9bb* | Proteobacteria | *Sphingomonadaceae* | -2.5 | 0.3 | 9.4 | 64 | 0 | 3.3E-18 |
| *c9aea435821b51653cfd1fdf3162bbab* | Gemmatimonadota | *Gemmatimonadaceae* | -2.4 | 0.2 | 10.0 | 58 | 0 | 8.1E-21 |
| *bafe930c8f8019dddab868bddc51f640* | Acidobacteriota | *Thermoanaerobaculia* | 1.7 | 0.3 | -6.3 | 54 | 59 | 1.3E-07 |
| *721bec93178db8ac44d9855db64fd88d* | Proteobacteria | *Reyranella soli* | 1.5 | 0.2 | -6.6 | 54 | 44 | 2.0E-08 |
| *26247a18506b8b2385fcb6460cac1500* | Actinobacteriota | *Solirubrobacter soli* | 2.4 | 0.2 | -15.4 | 49 | 107 | 1.2E-50 |
| *7c39ad081a428ca28a5cc21dd5c874bd* | Proteobacteria | *Mesorhizobium* | 1.9 | 0.3 | -5.8 | 45 | 71 | 5.0E-06 |
| *41b7521cb410f66d5fadd01ddfbc3d66* | Cyanobacteria | *Sericytochromatia* | 2.7 | 0.4 | -6.1 | 45 | 146 | 7.5E-07 |
| *f13a7640696f6cda15c6e1c42b844e7e* | Proteobacteria | *Sphingomicrobium* | -2.1 | 0.2 | 8.4 | 45 | 0 | 2.3E-14 |
| *ee49b2c458b7d8e54d794dac2230aaf1* | Proteobacteria | *Mesorhizobium* | 2.0 | 0.3 | -5.9 | 43 | 64 | 1.7E-06 |
| *77f48eb276896ac721d9660445ccd696* | Thermoproteota | *Nitrososphaeraceae* | 2.4 | 0.4 | -6.6 | 41 | 84 | 2.3E-08 |
| *2aaaf82eff5cd72a2d3cc69e5d4944a9* | Actinobacteriota | *Ilumatobacter coccineus* | -1.9 | 0.2 | 11.5 | 36 | 0 | 1.1E-27 |
| *6a3b81ace6a9a68bc16486f410c773e4* | Proteobacteria | *Xanthobacteraceae* | 2.1 | 0.3 | -7.9 | 34 | 56 | 1.3E-12 |
| *735994b04a3072f658972846a6f58909* | Acidobacteriota | *Pyrinomonadaceae* | -1.6 | 0.2 | 6.8 | 28 | 0 | 8.5E-09 |

| | | | | | | | | |
|---|---|---|---|---|---|---|---|---|
| *84bcc1f8b5e799a0b5fea9519520db9c* | Proteobacteria | *Sphingomonas* | -1.6 | 0.2 | 6.7 | 27 | 0 | 1.7E-08 |
| *b2af3ab5d31f6e0388d7e26fcb0b54d0* | Actinobacteriota | *Acidimicrobiales* | -1.5 | 0.3 | 5.0 | 27 | 0 | 2.7E-04 |
| *daac2813ec4defb7cdc75147d26333bc* | Verrucomicrobiota | *Verrucomicrobiaceae* | 3.1 | 0.6 | -4.8 | 19 | 50 | 9.7E-04 |
| *4d2718a6cb2774d782d35b0fb3f43e58* | Cyanobacteria | *Sericytochromatia* | 4.5 | 0.8 | -5.3 | 16 | 131 | 6.7E-05 |
| *0f995b24faa7fd071914de1aefaf880a* | Firmicutes_D | *Planococcaceae* | 3.9 | 0.7 | -5.3 | 14 | 63 | 8.6E-05 |
| *0669f23d503b4ff7ff013e65bc4ecb67* | Proteobacteria | *Micavibrio* | 3.4 | 0.7 | -5.1 | 6 | 25 | 1.8E-04 |
| *a6facc90b4326eef8109db25acb11d38* | Proteobacteria | *Xanthobacteraceae* | 3.8 | 0.8 | -4.8 | 0 | 17 | 8.1E-04 |
| *69f0863c21ffb4985a0586fdbfbd564c* | Proteobacteria | *Pseudolabrys* | 3.7 | 0.6 | -6.0 | 0 | 13 | 1.0E-06 |
| *4e893d68fbf03b0af9022a974dcb9265* | Planctomycetota | *Tepidisphaeraceae* | 4.2 | 0.9 | -4.8 | 0 | 26 | 8.9E-04 |
| *6870e333679b726bbb8993ea7b1c832f* | Acidobacteriota | *Vicinamibacteria* | 4.3 | 0.8 | -5.2 | 0 | 25 | 1.4E-04 |
| *2a13b4250e06e6fc3980788942bf86dc* | Proteobacteria | *Dongiaceae* | 3.9 | 0.7 | -5.2 | 0 | 16 | 1.3E-04 |
| *61f1dfd764ba55f5a494b3f5337141a8* | Proteobacteria | *Burkholderiaceae* | 3.7 | 0.8 | -4.9 | 0 | 13 | 5.8E-04 |
| *bb535db5a56df06eb50a972e6824b907* | Proteobacteria | *Usitatibacteraceae* | 4.0 | 0.7 | -6.1 | 0 | 18 | 5.8E-07 |

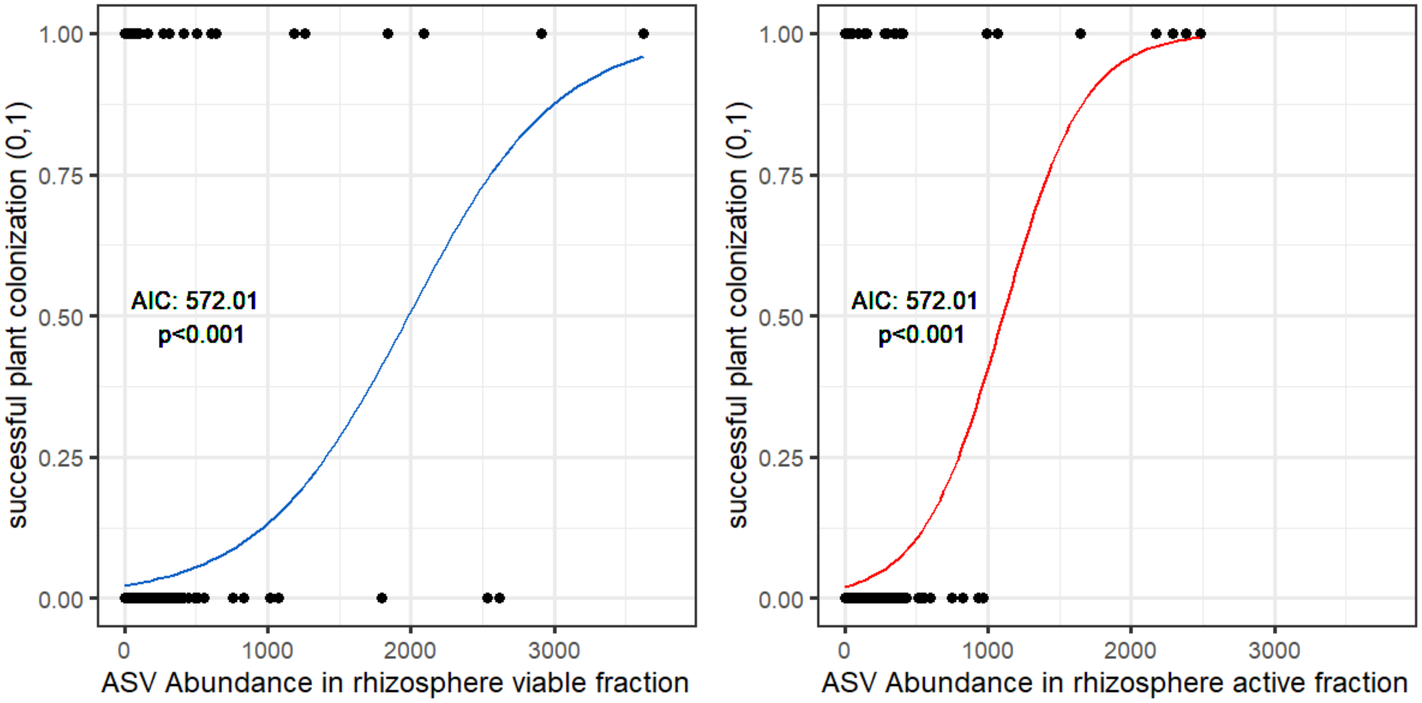


**Supplemental Figure 5: Successful plant colonization is more associated with abundance in the active fraction than viable fraction.** Plots visualize the binomial model of the association between ASVs abundance in the rhizosphere viable fraction (shown in blue) or in the rhizosphere active fraction (shown in red) with successful plant colonization. Successful plant colonization (binary) is defined by an ASV being present in the active or viable inside the roots or nodules with at least 50 reads. Model AICs are shown on the chart. P values are for abundance in the rhizosphere viable or active fraction. Both models used rarefied filtered datasets.

**Supplemental Table 8:** Results of binomial generalized linear model of successful plant colonization. Results from 1) null model 2) model ASVs abundance in the Viable fraction 3) ASVs abundance in the Active fraction. Estimates and Std. Error are untransformed log-odds.

**Null Model: presence in plant(1,0) ~ replicate plant**

**AIC: 600.32**

**Null deviance: 593.82 on 2467 degrees of freedom**

| Predictor | Estimate | Std. Error | Z value | P value |
|---|---|---|---|---|
| Intercept | -3.56E+00 | 2.46E-01 | -14.49 | <2e-16 |
| Rep2 | 5.88E-02 | 3.43E-01 | 0.171 | 0.864 |
| Rep3 | 1.17E-13 | 3.48E-01 | 0 | 1 |
| Rep4 | -3.57E-01 | 3.81E-01 | -0.935 | 0.35 |

**Model 2: presence in plant(1,0) ~ abundance in rhizosphere viable +replicate plant**

**AIC: 572.01**

**Null deviance: 593.82 on 2467 degrees of freedom**

| Predictor | Estimate | Std. Error | Z value | P value |
|---|---|---|---|---|
| Intercept | -3.68E+00 | 0.2482 | -14.85 | 2.00E-16 |
| Abundance Viable | 1.93E-03 | 0.0004 | 5.18 | 2.27E-07 |
| Rep2 | -1.05E-02 | 0.3512 | -0.03 | 0.976 |
| Rep3 | -7.20E-02 | 0.3545 | -0.20 | 0.839 |
| Rep4 | -4.30E-01 | 0.3900 | -1.10 | 0.27 |

**Model 3: presence in plant(1,0) ~ abundance in rhizosphere active**

**AIC: 555.46**

**Null deviance: 593.82 on 2467 degrees of freedom**

| Predictor | Estimate | Std. Error | Z value | P value |
|---|---|---|---|---|
| Intercept | -3.85E+00 | 0.2672 | -14.42 | 2.00E-16 |
| Abudance Active | 3.53E-03 | 0.0007 | 4.98 | 6.43E-07 |
| Rep2 | 1.06E-01 | 0.3598 | 0.30 | 0.767 |
| Rep3 | 3.49E-02 | 0.3650 | 0.10 | 0.924 |
| Rep4 | -3.59E-01 | 0.4013 | -0.89 | 0.371 |

**Supplementary Table 9:** Ten most abundant successful plant colonizers in the rhizosphere active fraction.These taxa contribute to the pattern in the binomial model.

| *ASVs* | *Phyla* | *Lowest taxonomic classification* | *Mean active* | *SD active* | *Mean viable* | *SD viable* |
|---|---|---|---|---|---|---|
| *f18c54e493d5d5ad534b9c3100216416* | *Proteobacteria* | *Rhizobium* | 2333.8 | 131.7 | 2451.5 | 1053.3 |
| *2acee427328d2e8641b8738258b15927* | *Proteobacteria* | *Pseudomonas* | 1022.3 | 516.7 | 1084.8 | 582.3 |
| *71582281edf705eb59a48c1a50e59a86* | *Proteobacteria* | *Devosia* | 364.5 | 10.6 | 326.5 | 43.1 |
| *509fca5a90d0b1dbec313e11c762a7e1* | *Proteobacteria* | *Rhizobium* | 347.0 | 43.7 | 347.3 | 149.3 |
| *e2b0e8123407c0522e65cc247a76628f* | *Proteobacteria* | *Rhizobiaceae* | 287.0 | NA | 56.0 | NA |
| *a799adf1c40ad36daa84625bd855e8ee* | *Proteobacteria* | *Variovorax paradoxus* | 249.0 | 44.7 | 188.0 | 31.4 |
| *3585021835f92062e3905fda1086a40f* | *Proteobacteria* | *Caulobacter* | 216.7 | 46.3 | 162.3 | 13.7 |
| *3e37cbf592eed8db558988be92d6cc49* | *Firmicutes_D* | *Neobacillus* | 189.5 | 181.7 | 70.5 | 40.3 |
| *1b86f9be5bce7a20bcf239e2b9d628b3* | *Proteobacteria* | *Caulobacter* | 173.0 | NA | 172.0 | NA |
| *d44ee1c36fcc9b098d56b04791f30386* | *Proteobacteria* | *Pseudomonas* | 156.5 | 89.7 | 165.3 | 83.0 |
